# ShinyAIM: Shiny-based Application of Interactive Manhattan Plots for Longitudinal Genome-Wide Association Studies

**DOI:** 10.1101/383026

**Authors:** Waseem Hussain, Malachy Campbell, Harkamal Walia, Gota Morota

**Author notes:** Corresponding author: Waseem Hussain, Department of Animal Science, Department of Agronomy and Horticulture, University of Nebraska-Lincoln, Lincoln, Nebraska 68583.

## Abstract

Due to advancements in sensor-based, non-destructive phenotyping platforms, researchers are increasingly collecting data with higher temporal resolution. These phenotypes collected over several time points are cataloged as longitudinal traits and used for genome-wide association studies (GWAS). Longitudinal GWAS typically yield a large number of output files, posing a significant challenge for data interpretation and visualization. Efficient, dynamic, and integrative data visualization tools are essential for the interpretation of longitudinal GWAS results for biologists but are not widely available to the community. We have developed a flexible and user-friendly Shiny-based online application, ShinyAIM, to dynamically view and interpret temporal GWAS results. The main features of the application include (i) an interactive Manhattan plots for single time points, (ii) a grid plot to view Manhattan plots for all time points simultaneously, (iii) dynamic scatter plots for p-value-filtered selected markers to investigate co-localized genomic regions across time points, (iv) and interactive phenotypic data visualization to capture variation and trends in phenotypes. The application is written entirely in the R language and can be used with limited programming experience. ShinyAIM is deployed online as a Shiny web server application at https://chikudaisei.shinyapps.io/shinyaim/, enabling easy access for users without installation. The application can also be launched on the local machine in RStudio.

## Introduction

Due to the increased availability of high-throughput phenotyping platforms, there is growing interest in the quantitative genetics of longitudinally measured traits, i.e., traits that are measured over multiple time points by advanced imaging systems (Araus and Kefauver 2018; Araus et al. 2018). For example, the application of GWAS to abiotic stress responses, such as drought, salinity, and temperature stress, measured at temporal resolution may provide insights into the mechanisms underlying plant physiological processes measured throughout the duration of stress or development (Busemeyer et al., 2013; Moore et al., 2013; Topp et al., 2013; Slovak et al., 2014; Wu◻rschum et al., 2014; Yang et al., 2014; Bac-Molenaar et al., 2015; Campbell et al 2015; Campbell, Walia, and Morota 2018).

Data visualization is a fundamental aspect of big data analysis in genetics. Manhattan plots are standard tools used to visualize GWAS results and to identify the genomic regions associated with a given phenotype. However, the static nature of these plots limits the information that can be displayed and extracted. Further, the number of Manhattan plots that can be viewed at one time is limited, making comparisons across phenotypes tedious. The situation becomes more challenging in the case of longitudinal GWAS, which are performed across multiple time points, with each time point producing a Manhattan plot. Furthermore, it is difficult to share GWAS outputs in an easy and convenient way, requiring novel applications for dynamic data visualization and sharing. Many browsers have been built to visualize GWAS outputs (e.g., Khramtsova and Stranger 2017; Cuellar-Partida, Renteria, and MacGregor 2015; Juliusdottir et al. 2018; Ziegler, Hartsock, and Baxter 2015). However, none of these are specifically tailored for longitudinal traits. Further, existing applications do not offer features for the dynamic visualization of Manhattan plots online and for comparisons across timepoints simultaneously.

To address these limitations, we have developed a Shiny-based application, ShinyAIM, for visualizing and interpreting longitudinal GWAS outputs in an interactive way. The application is distinct from previously developed GWAS visualization browsers because it is specifically designed for longitudinal traits, allowing the simultaneous visualization of all time points or phenotypes and comparisons of top associated markers across time points. The interactive and integrative GWAS and phenotypic data visualization features embedded in the application offer a new resource for users to readily extract extensive information from temporal GWAS results.

## Overview of ShinyAIM

### Methods

ShinyAIM is entirely written in the R language (R Core Team 2018) with the underlying R code encapsulated by the shiny R package (Chang et al. 2018), which is a web application framework for R, offering an interactive graphical user interface. Shiny has been making inroads into plant breeding and quantitative genetics for research and teaching purposes, such as Be-Breeder (Fritsche-Neto and Matias, 2016) and ShinyGPAS (Morota 2017). ShinyAIM leverages the cumulative utility of the R packages manhattanly (Sahir 2016), plotly (Sievert et al. 2017) and ggplot2 (Wickham 2016) to create a cohesive web browser-based application. The ShinyAIM application does not require any working knowledge of R and is intuitively operated through graphical user interface. ShinyAIM is hosted by a Shiny web server (https://chikudaisei.shinyapps.io/shinyaim/) for online use or can be run locally within RStudio by running the code *shiny::runGitHub(“ShinyAIM”, “whussain2”*). Alternatively, the ShinyAIM source code and sample files can be directly downloaded from the GitHub repository (https://github.com/whussain2/ShinyAIM). From the downloaded directory, the source file named app.R in RStudio can be run by clicking the *Run App* button. The ShinyAIM application is open source and is distributed under Artistic License 2.0.

### Usage

The starting page of the ShinyAIM application includes the Information tab with detailed information on how to format and upload the data. The video demonstration illustrating the application usage is also available (https://youtu.be/5-JLMpSiwv4). The ShinyAIM is aimed for visualization of GWAS outputs and does not perform GWAS analysis. There are five required columns in the user data file labeled as ‘timepoint’ (time point), ‘marker’ (marker name), ‘chrom’ (chromosome number), ‘pos’ (marker position), and ‘P’ (marker p-value) for Manhattan plot visualizations. For phenotypic data visualization, the data file must have two columns including ‘timepoint’ (time point), and ‘Value’ (phenotypic value). Further detailed instructions regarding the data formatting and column naming can be found in the main Information tab. In addition, the sample data files can be directly downloaded by clicking the ‘Download Sample File’ button given on the top of sidebar panel in the main tab.

The ShinyAIM application hosted on the server can handle 200-300k markers for the visualization of interactive Manhattan plots. However, we suggest to launch the application locally by running the code *shiny::runGitHub(“ShinyAIM”, “whussain2”)* in RStudio for the datasets with millions of markers. Alternatively, filtering can be done based on p-values by removing markers with large p-values prior to uploading the input file for visualization.

### Main features and functionality

The application has four main features to explore GWAS results: (i) interactive Manhattan plots for single time points, (ii) Manhattan grid plot to compare results across all time points simultaneously, (iii) dynamic views of p-value filtered top associated markers in a scatter plot to identify co-localized markers over time, and (iv) visualization of phenotypic data used for GWAS (Figure 1). These features are supported by user-defined data filtering criteria in ShinyAIM to smoothly navigate the application. Each feature is briefly described in the following sections.

**Figure 1:**
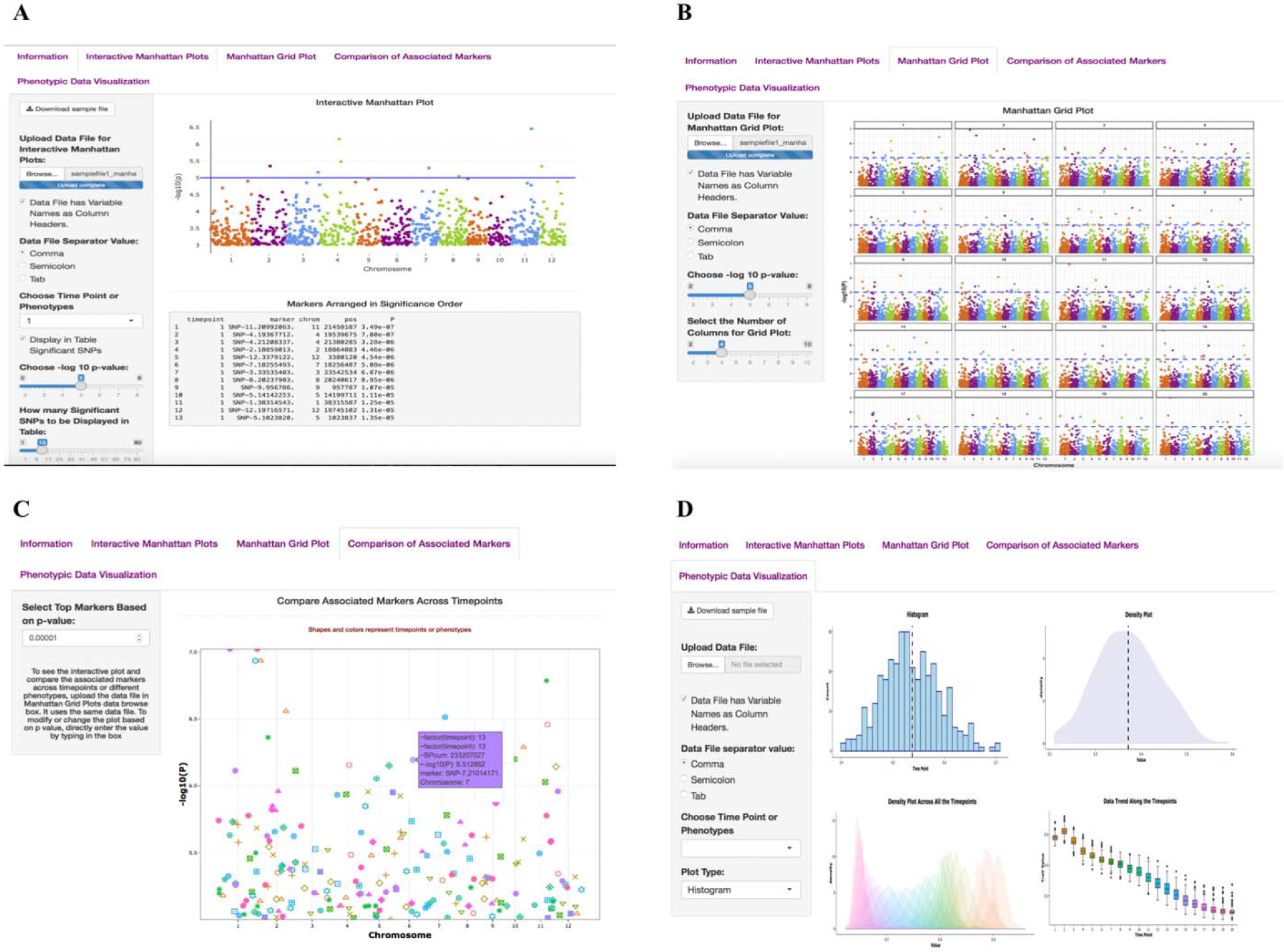
Main interface of the ShinyAIM application. Screenshots of panels for the main tabs are shown. (A) The ‘Interactive Manhattan Plots’ tab allows users to display interactive Manhattan plots for a selected time point. Users have the flexibility to choose the significance level and can display the top associated markers in tabular format. (B) The ‘Manhattan Grid Plot’ tab allows users to visualize Manhattan plots for all time points simultaneously. Users have the flexibility to choose the significance level and the number of columns in the grid plot. (C) The ‘Comparison of Associated Markers’ tab allows users to filter markers based on p-values, display a scatter plot for comparisons across all time points, and search for co-localized markers. (D) The ‘Phenotypic Data Visualization’ tab generates histogram and density plots and summarizes trends in temporal phenotypic data in the form of box plots.

### Interactive Manhattan Plots

In the Interactive Manhattan Plots panel, users can interactively view the Manhattan plot for each time point (Figure 1A). After the correct file format is selected and the file is uploaded, the available time points will be automatically updated in the ‘Choose Time Point or Phenotypes’ menu. An interactive Manhattan plot is automatically generated on the right-hand panel after selecting a target time point. Users can move the mouse over the points in the plot to display detailed information, including the marker name, position, chromosome location, and −log10 p-value. Furthermore, it is possible to zoom in on potential candidate regions to obtain additional detail. ShinyAIM offers the flexibility to choose the significance level by moving the slider input bar. In addition, users have a choice to display a list of markers arranged in decreasing order of p-values in the table below the Manhattan plot panel. The display also includes marker information in the input data file. The slider input bar controls the number of markers shown in the table.

### Manhattan Grid Plot

Manhattan Grid Plot tab allows users to visualize the Manhattan plots combined for all time points and can be used to explore how GWAS peaks change over time to facilitate data interpretation (Figure 1B). The significance threshold for markers can be modified by moving the slider input bar. Moreover, ShinyAIM enables users to choose the number of columns and rows in the grid plot by moving the slider input bar ‘Select the Number of Columns in Grid Plot.’

### Comparison of Associated Markers

Users are able to dynamically view only the top associated markers in a scatter plot (Figure 1C). This feature is implemented in ShinyAIM to enable users to focus only on the topmost associated markers and compare these markers across time points to identify co-localized regions. Users can select the number of markers displayed in a scatter plot by filtering the markers based on p-values. This is achieved by directly typing or selecting the option ‘Select Top Markers Based on p-value.' The scatter plot is interactive, and users can move the mouse over a point to display information, including the time point, chromosome name, position of the marker, name of the marker, and −log10 p-value (Figure 1C).

### Phenotypic Data Visualization

Phenotypic data visualization helps users view phenotypes used for GWAS in the form of dynamic histograms and density plots (Figure 1D). The trends and variability in phenotypic values at each time point can be visualized using box plots. All plot types are interactive, and users can move the mouse over a particular point to obtain detailed information.

### Conclusion

We have developed a user-friendly integrative Shiny-based application to dynamically visualize and interpret longitudinal GWAS results, providing an easy-to-use online tool to the community.

### Availability

The source code for the ShinyAIM application is freely available at the GitHub repository https://github.com/whussain2/ShinyAIM or at the Zenodo repository https://zenodo.org/record/1422835. The source code is licensed under Artistic License 2.0. ShinyAIM can be launched on any system that has RStudio installed or available online at the Shiny web server https://chikudaisei.shinyapps.io/shinyaim/

## Acknowledgments

This work was funded and supported by National Science Foundation Grant No.1736192.

## References

Araus, J. L., & Kefauver, S. C. (2018). Breeding to adapt agriculture to climate change: Affordable phenotyping solutions. Current Opinion in Plant Biology (In Press). https://doi.org/10.1016/j.pbi.2018.05.003

Araus, J.L., Kefauver, S. C., Zaman-Allah, M., Olsen, M. S., & Cairns, J. E. (2018). Translating high-throughput phenotyping into genetic gain. Trends in Plant Science, 23(5), 451–466. https://doi.org/10.1016/j.tplants.2018.02.001.

Bac-Molenaar, J. A., Vreugdenhil, D., Granier, C., & Keurentjes, J. J. B. (2015). Genome-wide association mapping of growth dynamics detects time-specific and general quantitative trait loci. Journal of Experimental Botany, 66(18), 5567–5580. http://dx.doi.org/10.1093/jxb/erv176

Bhatnagar, S. (2016). Manhattanly: Interactive Q-Q and Manhattan Plots Using “Plotly.Js.” https://CRAN.R-project.org/package=manhattanly.

Busemeyer, L., Ruckelshausen, A., Möller, K., Melchinger, A. E., Alheit, K. V., Maurer, H. P, Hahn, V., Weissmann, E. A., Reif, J. C., & Wu◻rschum, T. (2013). Precision phenotyping of biomass accumulation in triticale reveals temporal genetic patterns of regulation. Scientific Reports, 3, 2442. https://www.nature.com/articles/srep02442

Campbell, M. T., Walia, H., & Morota, G. (2018). Utilizing random regression models for genomic prediction of a longitudinal trait derived from high-throughput phenotyping. BioRxiv 319897 [Preprint]. https://doi.org/10.1101/319897

Chang, Winston., Cheng, J., & Allaire, et al. (2018) Shiny: Web application framework for R. https://CRAN.R-project.org/package=shiny.

Cuellar-Partida, G., Renteria, M. E., & MacGregor, S. (2015). LocusTrack: Integrated Visualization of GWAS Results and Genomic Annotation. Source Code for Biology and Medicine, 10, 1. https://doi.org/10.1186/s13029-015-0032-8

Fritsche-Neto, Roberto., & Matias, F. I. (2016). Be-Breeder - Learning: A new tool for teaching and learning plant breeding principles. Crop Breeding and Applied Biotechnology, 16(3), 240–245. http://dx.doi.org/10.1590/1984-70332016v16n3n36.

Wickham, H. (2016). ggplot2: Elegant Graphics for Data Analysis. Springer-Verlag New York. ISBN 978-3-319-24277-4. http://ggplot2.org.

Juliusdottir, T., Banasik, K., Robertson, N. R., Mott, R., & McCarthy, M. I. (2018). Toppar: An Interactive Browser for Viewing Association Study Results. Bioinformatics, 34(11), 1922–1924. https://doi.org/10.1093/bioinformatics/btx840

Khramtsova, E. A., & Stranger, B. E. (2017). Assocplots: A Python package for static and interactive visualization of multiple-group GWAS results. Bioinformatics, 33(3), 432–434. https://doi.org/10.1093/bioinformatics/btw641

Moore, C. R., Johnson, L. S., Kwak, I. Y., Livny, M., Broman, K. W., & Spalding, E. P. (2013). High-throughput computer vision introduces the time axis to a quantitative trait map of a plant growth response. Genetics, 195, 1077–1086. https://doi.org/10.1534/genetics.113.153346

Morota, G. (2017). ShinyGPAS: Interactive Genomic Prediction Accuracy Simulator based on deterministic formulas. Genetics, Selection, Evolution 49: 91. https://doi.org/10.1186/s12711-017-0368-4

R Core Team. (2018). R: The R Project for Statistical Computing. Vienna, Austria. https://www.r-project.org/.

Sievert, C., Parmer, C., Hocking, et al. (2017). Plotly: Create Interactive Web Graphics via “Plotly.Js.” https://CRAN.R-project.org/package=plotly

Slovak, R., Göschl, C., Su, X., Shimotani, K., Shiina, T., & Busch, W. (2014). A scalable open-source pipeline for large-scale root phenotyping of Arabidopsis. Plant Cell, 26, 2390–2403. https://doi.org/10.1105/tpc.114.124032

Topp, C. N., Iyer-Pascuzzi, A. S., Anderson, J. T., Lee, C. R., Zurek, P. R., Symonova, O., Zheng, Y., Bucksch, A., Mileyko, Y., Galkovskyi, T., et al. (2013). 3D phenotyping and quantitative trait locus mapping identify core regions of the rice genome controlling root architecture. Proceedings of National Academy of Science USA, 110: E1695–E1704. https://doi.org/10.1073/pnas.1304354110

Wu◻rschum, T., Liu, W., Busemeyer, L., Tucker, M. R., Reif, J.C., Weissmann, E. A., Hahn, V., Ruckelshausen, A., & Maurer, H. P. (2014). Mapping dynamic QTL for plant height in triticale. BMC Genetics, 15, 59. https://doi.org/10.1186/1471-2156-15-59

Yang, W., Guo, Z., Huang, C., Duan, L., Chen, G., Jiang, N., Fang, W., Feng, H., Xie, W., Lian, X., et al. (2014). Combining high-throughput phenotyping and genome-wide association studies to reveal natural genetic variation in rice. Nature Communications, 5, 5087. https://www.nature.com/articles/ncomms6087

Ziegler, G. R., Ryan, H. H., & Baxter, I. (2015). Zbrowse: An interactive GWAS results browser. PeerJ Computer Science 1, e3. https://peerj.com/articles/cs-3/

